# Cholinergic interneurons in the nucleus accumbens are a site of cellular convergence for corticotropin release factor and estrogen regulation

**DOI:** 10.1101/2024.04.13.589360

**Authors:** Kendra Olson, Anna E. Ingebretson, Eleftheria Vogiatzoglou, Paul G. Mermelstein, Julia C. Lemos

## Abstract

Cholinergic interneurons (ChIs) act as master regulators of striatal output, finely tuning neurotransmission to control motivated behaviors. ChIs are a cellular target of many peptide and hormonal neuromodulators, including corticotropin releasing factor, opioids, insulin and leptin, which can influence an animal’s behavior by signaling stress, pleasure, pain and nutritional status. However, little is known about how sex hormones via estrogen receptors influence the function of these other neuromodulators. Here, we performed in situ hybridization on mouse striatal tissue to characterize the effect of sex and sex hormones on choline acetyltransferase (*Chat*), estrogen receptor alpha (*Esr1*), and corticotropin releasing factor type 1 receptor (*Crhr1*) expression. Although we did not detect sex differences in ChAT protein levels in the striatum, we found that female mice have more *Chat* mRNA-expressing neurons than males. At the population level, we observed a sexually dimorphic distribution of *Esr1*- and *Crhr1*-expressing ChIs in the ventral striatum that demonstrates an antagonistic correlational relationship, which is abolished by ovariectomy. Only in the NAc did we find a significant population of ChIs that co-express *Crhr1* and *Esr1*. At the cellular level, *Crhr1* and *Esr1* transcript levels were negatively correlated only during estrus, indicating that changes in sex hormones levels can modulate the interaction between *Crhr1* and *Esr1* mRNA levels. Together, these data provide evidence for the unique expression and interaction of *Esr1* and *Crhr1* in ventral striatal ChIs, warranting further investigation into how these transcriptomic patterns might underlie important functions for ChIs at the intersection of stress and reproductive behaviors.

## Introduction

The hypothalamus-pituitary-“peripheral organ” systems and cortico-striatal-mesolimbic system both play important roles in maintaining homeostasis. Broadly speaking, the hypothalamus detects homeostatic disturbances or changes across several internal or external dimensions including changes in caloric intake/blood glucose levels and water/osmotic levels, introduction of threats in the environment and presentation of salient reproductive cues. Motivational circuits that encompass the nucleus accumbens (NAc) and ventral tegmental area (VTA) are critical in driving actions that allow the animal to essentially rectify or address these homeostatic disturbances from acquisition of food to defense of a territory^1–6^. As such, these peptides and hormones with their accompanying receptors present in the hypothalamus are also present in the cortico-striatal-mesolimbic pathway. Within the hypothalamus, it has been shown the sexual dimorphism and circulating sex hormones effect the function other hormonal systems associated with stress, feeding and pain to modulate homeostatic sensitivity and behavioral output^7–9^. Yet the precise way that these neuropeptides and hormones regulate this cortico-striatal-mesolimbic pathway and interact with each other is understudied.

Cholinergic interneurons (ChIs) in the dorsal and ventral striatum are the main source of striatal acetylcholine^10–12^. Despite only constituting 1-2% of all neurons in the striatum, their morphological and functional attributes position them to act as master regulators or “gatekeepers” of striatal output^12,13^. Acetylcholine transmission, via ionotropic nicotinic acetylcholine receptors (nAChRs) and metabotropic muscarinic acetylcholine receptors (mAChRs), modulates the excitability and fast synaptic transmission of the two major classes of GABAergic projection neurons: direct, or dopamine D1 receptor-containing spiny projection neurons, and indirect, or dopamine D2 receptor-containing spiny projection neurons^14–16^. In addition, ChIs have been shown to modulate dopamine transmission through the activation of either mAChRs^17–22^ or nAChRs, with nAChR activation on dopaminergic varicosities capable of generating action potential-mediated dopamine release that is independent of somatic firing^17–22^. This relationship between acetylcholine and dopamine transmission is critical to maintaining striatal function.

Cholinergic and dopaminergic neurons in the striatum also have unique and interacting signatures of activity during reward learning^23–25^, and it has been shown that reducing ChI activity, specifically in the nucleus accumbens (NAc), produces anxiety- and depression-like phenotypes in mice^26–28^. A recent study by Brady et al. demonstrated that nAChR-triggered dopamine transmission is different between males and females, suggesting that there may be fundamental sex differences in the relationship between striatal acetylcholine and dopamine^29^. This finding may explain the sex biases observed in several striatal-dependent neuropsychiatric disorders, but the influence of sex on striatal function is not fully understood. Therefore, given the importance of ChIs in controlling the network function and output of the striatum, our goal was to answer fundamental questions regarding potential sex differences in the number and distribution of ChIs across subregions of the striatum.

ChIs are a cellular target of many peptide and hormonal neuromodulators. Opioids, insulin, leptin and corticotropin releasing factor (CRF) can each modulate the firing of ChIs and can therefore indirectly modulate dopamine transmission in the striatum^19,30–36^. We have previously shown, in male mice, that the vast majority (80-90%) of ChIs express CRF type 1 receptors (CRF-R1), that CRF increases the firing of ChIs and that this increases dopamine transmission via activation of muscarinic M5 receptors located on dopaminergic *en passant* terminals^19^. While ongoing studies are assessing this electrophysiological effect in female mice, we first wanted to examine the localization of CRF-R1 receptors relative to estrogen receptors in female mice to determine if there is a direct interaction between estrogen receptors and CRF receptors at the site of ChIs.

Across species, female sex hormones fluctuate across the reproductive cycle. Across species, female sex hormones fluctuate across the reproductive cycle. Mice have approximately a four-day cycle where estradiol and progesterone peak across proestrus, driving sexual motivation during estrous, and return to low levels during metestrus and diestrus^37^. Across the estrous cycle, changes in estrogen receptors themselves have been documented in areas of the hypothalamus^38,39^. Importantly, it has been shown that even in this short time span, structural and functional plasticity occurs within these motivational circuits^40–42^. Estradiol can influence neuronal function via both nuclear and/or surface membrane localized receptors^43–47^. Canonically, when bound to estradiol, the estrogen receptors, ERα and ERβ bind DNA to regulate gene expression^43,44^. Through palmitoylation, these same estrogen receptors are trafficked to the cell membrane where they induce changes in excitability and synaptic transmission by functionally coupling to metabotropic glutamate receptors^46^. It has been shown that acute application of estradiol can rapidly decrease glutamatergic synaptic transmission, particularly in the NAc of intact females^48,49^. Indeed, there appears to be profound regional differences in the effect of estradiol or hormonal cycle on synaptic transmission, with no significant effects in the dorsal striatum. It has also been shown that glutamatergic transmission can be impacted by estrous cycle. This has largely been attributed to ERα (gene name *Esr1*). The effects of sex and sex hormones can be disrupted by removing the ovaries of female animals, an effect that not only shuts down circulating sex hormone production but also impairs the normal function of stress-associated hormones like corticosterone^50^.

Electron microscopy studies examining ERα have shown limited expression in the striatum. ERα is not expressed heavily in spiny projection neurons post-synaptically; rather its presence has been noted on astrocytes, ChIs and other GABAergic interneurons within the NAc — though sex dependent differences in expression profiles have not been assessed^51,52^. Given the known behavioral interaction of sex and sex steroid cycle with stress, feeding behaviors and pain sensitivity^7–9^, we speculated that one potential role of striatal ERα is to regulate the expression and function of other GPCRs such as CRF-R1s, especially in theses “gatekeepers” of striatal output: the ChIs. We therefore assessed sex- and sex-steroid dependent changes in ERα and CRF-R1 mRNA expression across regions of the striatum, both overall and at the level of ChIs using fluorescent *in situ* hybridization strategies.

## Methods

### Animals

All procedures were done in accordance with the University of Minnesota Animal Care and Use Committee. Both male and female C57BL6/J mice were used in these studies at ages post-natal day 60-160. Mice were group-housed and kept under a 12h light cycle (6:00 ON/18:00 OFF). Ovariectomized C57BL6/J females were purchased from Jackson Laboratories. Food and water were available *ad libitum*.

### 17β-estradiol (E2) administration

Ovariectomized (OVX) females were treated with a single administration of 17β-estradiol (E2, Sigma-Aldrich cat# 3301; dissolved in cottonseed oil to a concentration of 2 µg/0.1mL and injected subcutaneously at a volume of 0.1 mL) or vehicle 5 or 72 hours prior to tissue collection. We did not see any differences in *Crhr1* or *Esr1* mRNA expression after 5 or 72 hours of E2 treatment (*Crhr1* expression unpaired t-test p = 0.2756, 5 hours: N = 4, 72 hours: N = 4; *Esr1* expression unpaired t-test p = 0.8689, 5 hours: N = 4, 72 hours: N = 4), therefore these data were pooled.

### Estrous cycle tracking

Vaginal lavage methods were used to track the estrous cycle of female mice over an 8–10 day period in order to capture two full cycles. Briefly, 20 µl of sterile saline was rapidly pipetted in and out of the vagina such that vaginal cells would be collected within the solution. The sample was dried, stained with cresyl violet, and assessed under the microscope. Females were classified as being in proestrus, estrus, metestrus or diestrus based on the proportion of leukocytes, cornified epithelial cells and nucleated epithelial cells, as well as the size of the vaginal opening. We did not find any sex differences or effect of estrous cycle between the NAc core versus NAc shell in *Crhr1* or *Esr1* expression overall or in ChIs specifically (data not shown), and therefore data from these two regions were pooled.

### Fluorescent in situ hybridization (RNAscope, ACD)

Mice were anesthetized with isoflurane and rapidly decapitated. Brains were extracted and flash-frozen in isopentane on dry ice. Brains were stored at −80°C until sectioning. Prior to sectioning, brains were moved to −20°C and acclimated for 2–24 hours. Using a Leica CM8060 cryostat, 16µm slices were thaw mounted onto slides (Superfrost, Fisher) and stored at −80°C until assayed. RNAscope manual fluorescent multiplex v1 were run as detailed in ACD technical manuals and previously reported (see https://acdbio.com/main-category/user-manuals). For all *in situ* hybridization experiments, N represents animals except in Figure 5i-l where N represents individual cells. For sex comparisons, a total of 20-24 unique images per animal were acquired. The following probes were purchased from ACD: Mm-Crhr1-C1 (Cat#: 418011), Mm-Esr1-C2 (Cat #: 478201-C2) and Mm-Chat-C3 (Cat #: 408731-C3).

### Immunohistochemistry

Mice were intracardially perfused with phosphate-buffered saline (PBS) containing 10U/mL Heparin until the liver cleared (approximately 5 minutes) followed by 40 mL fixative solution (4% paraformaldehyde; 4% sucrose in 0.1M PBS) at 3 mL/min. Brains were incubated in 30% sucrose (in 0.1 M PBS) for 24 hours at 4°C and sliced with a cryostat or microtome. 40µm floating sections were washed in PBS and then blocked for 1 hour in 5% donkey serum, 0.3% Triton-X in PBS at room temperature (RT). Sections were then incubated in primary antibody for 12-18 hours at RT (goat anti-ChAT, 1:500, Millipore Sigma and Cat#: AB144P; mouse anti-NeuN Clone A60-AlexFluor 488 conjugated, Millipore Sigma and Cat #: MAB377X). Slices were incubated in donkey anti-goat Alexa Fluor 555 (1:500, Invitrogen) for 2 hours at RT. Slices were washed 3 times (10 min each) in PBS, mounted with Vectashield hard set mounting media containing DAPI (Vector Laboratories and Cat #:H-1500), and coverslipped.

### Imaging and analysis

Images (1024×1024 pixels) were acquired using a confocal microscope (Stellaris 8 or AR-FLIM NIKON Confocal microscope). Six to eight confocal images were acquired in each region per mouse. Analysis was performed using ImageJ or Indica Halo software on acquired images. For RNAscope experiments, “single plane” (5µm thickness) 40x confocal images of *Crhr1, Chat* and *Esr1* mRNA expression in the dorsal striatum, NAc core, and NAc shell were acquired. ImageJ was used to set a constant threshold in order to analyze both total numbers of positive and negative cells and also to count number of puncta. Different mRNA probes in the ACD catalog have differing sensitivities. Thus, for a unique mRNA probe, the threshold for minimum intensity, puncta size and number of puncta was titrated based on signal quality and expression pattern of each mRNA probe (i.e. *Crhr1* vs. *Esr1* vs. *Chat*). However, the same imaging and analysis settings were applied across all sections and all animals from a single RNAscope run for each probe. Settings were adjusted as needed between runs, and similar numbers of animals from different experimental groups were used for each run when possible. A threshold of 5 puncta was set to consider a cell positive for mRNA expression, with number of puncta being an approximation of total number of transcripts. For analysis of immunohistochemistry, the HALO image analysis platform (Indica Labs) was used. The dorsomedial striatum (DMS), dorsolateral striatum (DLS), NAc core, and NAc shell were identified during image collection. In this study, four 20 µm z-stack at 40x magnification was collected for each region, and the maximum

### Statistics

Statistical analysis was performed in Prism (GraphPad) and Excel. Many analyses used a one-way ANOVAs with sex or estrous cycle as the independent variable. In some cases, a two-way ANOVA was used with region and sex as the two factors in the analysis. When stated, unpaired t-tests were used. For correlational analysis, a Pearson’s correlation was run. Data are reported as mean ± SEM.

## Results

### Sex-dependent differences in ChAT mRNA expression, but not protein expression

We first aimed to determine if there were any differences in ChI number or density in different striatal subregions of male and female mice. ChIs are reliably identified by the expression of choline acetyltransferase mRNA and/or protein, the rate limiting enzyme for acetylcholine production^53^. To test this, we used both immunohistochemistry and *in situ* hybridization to assess differences in protein expression and mRNA, respectively. While immunohistochemistry has the benefits of detecting protein expression, it generally has lower sensitivity than measuring gene or RNA expression. We conducted standard immunohistochemistry procedures in male and female mice assessing ChAT-immunoreactivity (ChAT-ir) in the dorsolateral striatum (DLS), dorsomedial striatum (DMS), NAc core and NAc shell. We counterstained for NeuN (a neuronal marker) so that we could assess total number of ChAT-ir+ cells as well as the density of ChAT-ir+ cells relative to total neurons. Unlike many antibodies for GPCRs, the ChAT and NeuN antibodies used are well-validated and reliable. When assessing both measures, we replicated our previously published regional differences in ChAT-ir+ cells in which we see fewer ChAT-ir+ cells in the NAc core compared to DLS, DMS or NAc shell (two-way ANOVA, main effect of region, total # ChAT-ir+: F_3,20_ = 8.490, p = 0.008; % co-localized with NeuN: F_3,20_ = 10.66, p = 0.0002, N = 3 male, 4 female, Figure 1a-c). We did not find any significant main effect of sex or sex by region interaction in either the total number of ChAT-ir+ cells (main effect of sex, p = 0.1869; sex by region interaction, p = 08170) or percent of ChAT-ir+ cells that were colocalized with NeuN (main effect of sex, p = 0.1427, sex by region interaction, p = 0.7177).

**Figure 1.**
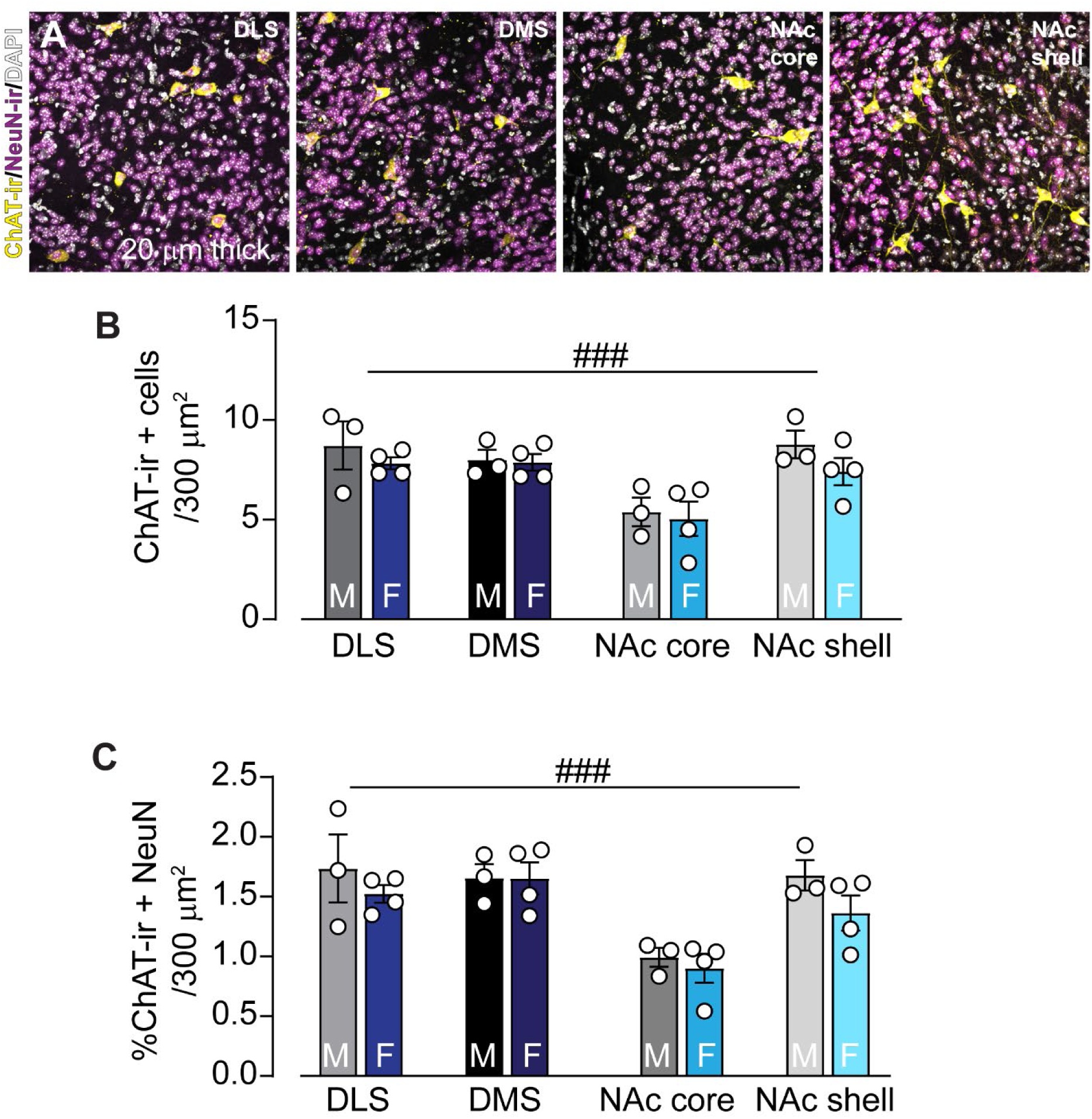
Region and sex-dependent differences in ChAT-immunoreactivity levels. A) Example images of ChAT-immunoreactivity (ir; yellow) is seen in neurons (NeuN, magenta) throughout the dorsolateral striatum (DLS), dorsomedial striatum (DMS), nucleus accumbens (NAc) core, and NAc shell. Nuclei indicated with DAPI in grey. Images are maximum projections of 20 µm thick z-stacks. B) Fewer ChAT-ir cells are located in the NAc core compared to the DLS, DMS, or NAc shell (two-way ANOVA, main effect of region, F_3,20_ = 8.490, p = 0.008), with no differences in number of ChAT-ir cells between males (M, gray, N = 3) and females (F, blue, N = 4; p = 0.1869). C) Percent of ChAT-ir cells that colocalized with the neuronal NeuN marker. ChAT-ir neurons are less dense in the NAc core compared to the DLS, DMS, or NAc shell (two-way ANOVA, main effect of region, F_3,20_ = 10.66, p = 0.0002) with no effect of sex (p = 0.1427). ###: two-way ANOVA, main effect of region, p < 0.01.

We next analyzed *Chat* mRNA expression using *in situ* hybridization, and compared overall numbers of *Chat*+ cells as well as overlap with DAPI (a nucleus marker) to determine the density of *Chat*+ cells relative to total nuclei. Here, we replicated the finding that there were more *Chat*+ cells in the dorsal striatum than in the NAc, as expected (two-way ANOVA, main effect of region, total # *Chat*+: F_1,40_ = 18.11, p = 0.0001; % co-localized with DAPI: F_1,40_ = 5.839, p = 0.0203, N = 6-26, Figure 2a,b). However, in contrast to our protein analysis where no sex differences were found, here we found that females had a small, but significant elevation in the total number of *Chat* mRNA-expressing cells in both the dorsal striatum and NAc. This was also true when we analyzed the density of *Chat+* cells co-localized with DAPI (two-way ANOVA, main effect of sex, total #: F_1,40_ = 4.666, p = 0.0368; % co-localized with DAPI: F_1,40_ = 29.26, p < 0.0001, N = 6-26, Figure 2a,c). These data indicate that there may be a sex-dependent change in the striatal ChAT mRNA to ChAT protein ratio. Alternatively, it is possible that the *in situ* hybridization method is more sensitive than the immunohistochemistry method.

**Figure 2.**
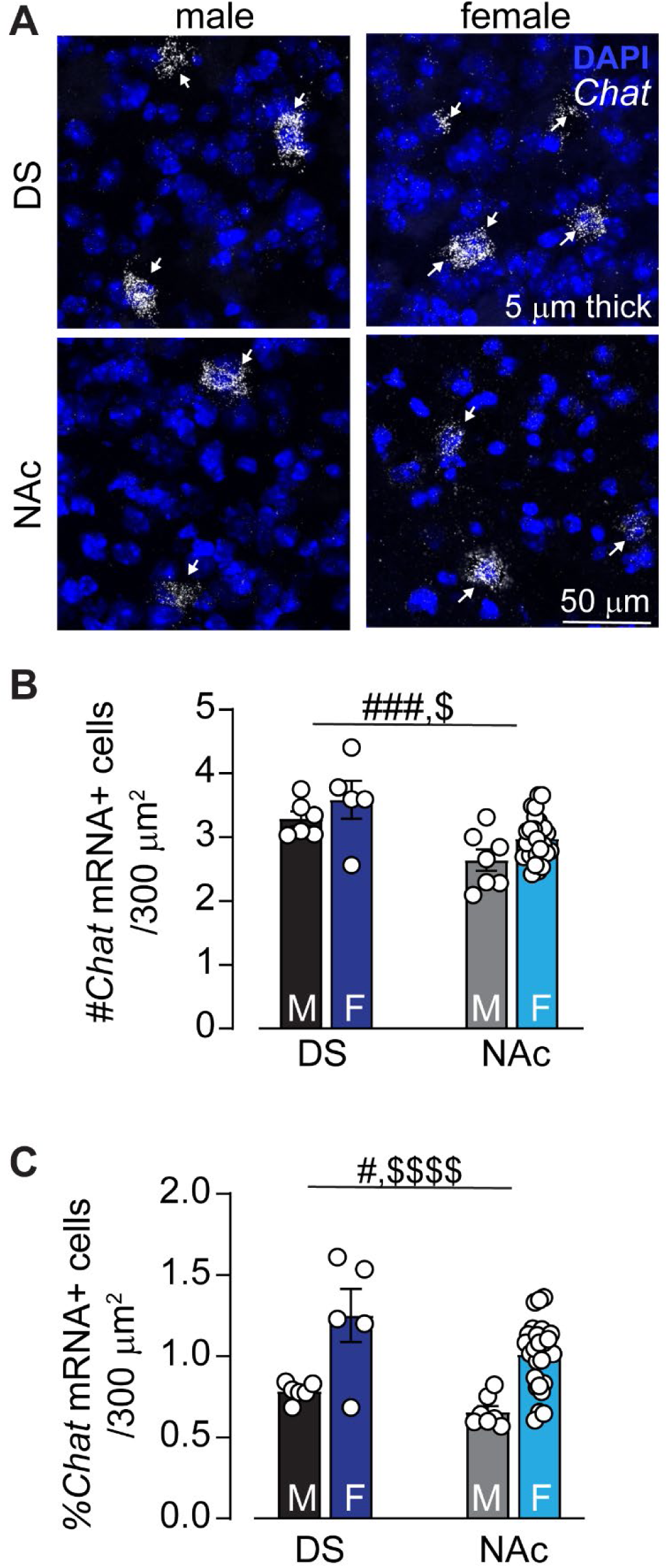
Sex differences in *Chat* mRNA expression. A) Example images of *Chat* mRNA visualized in the dorsal striatum (DS) and nucleus accumbens (NAc) using *in situ* hybridization alongside DAPI for nucleus visualization. Images are from 5µm thick “single plane” from an example male and female mouse. B) Fewer *Chat* mRNA+ cells are located in the NAc compared to the DS (two-way ANOVA, main effect of region, F_1,40_ = 18.11, p = 0.0001, N = 6-26), with females having more *Chat*+ cells in both the NAc and DS compared to males (two-way ANOVA, main effect of sex, F_1,40_ = 4.666, p = 0.0368, N = 6-26). C) Percent of *Chat*+ cells co-localized with DAPI. *Chat+* cells are less dense in the NAc compared to the DS (two-way ANOVA, main effect of region, F_1,40_ = 5.839, p = 0.0203), with females having notably denser *Chat*+ cells in both the NAc and DS compared to males (two-way ANOVA, main effect of sex, F_1,40_ = 29.26, p < 0.0001, N = 6-26). #: Two-way ANOVA main effect of region p < 0.05. ###: Two-way ANOVA main effect of region p < 0.01. $: Two-way ANOVA main effect of sex p < 0.05. $$$$: Two-way ANOVA main effect of sex p < 0.0001.

### No sex differences in Crhr1 and Esr1 in the dorsal striatum

Previous work from our lab indicates that *Crhr1* is robustly expressed on ChIs in the dorsal and ventral striatum, strongly regulating their firing. Since there are clear interactions between sex and stress hormonal systems in the hypothalamus, we hypothesized that there may be sex- and estrous-cycle differences in *Esr1* and *Crhr1* expression overall, and in ChIs specifically, in the dorsal and ventral striatum. To test this, we used *in situ* hybridization techniques.

There was no significant difference in the density of *Crhr1+* cells in the dorsal striatum between males (8.1±0.5%) and females (7.2±0.9%) (unpaired t-test, t = 0.9380, p = 0.3703, N = 6 males, 6 females, Figure 3a,b). The expression of *Esr1*+ cells overall was very sparse in the dorsal striatum (0.44±0.16% for males and 0.35±0.05% for females) and also did not differ between males and females (unpaired t-test, t = 0.5742, p = 0.5785, Figure 3a,c). These data suggest there are no sex differences in the density of *Esr1*+ and *Crhr1*+ cells in the mouse dorsal striatum.

**Figure 3.**
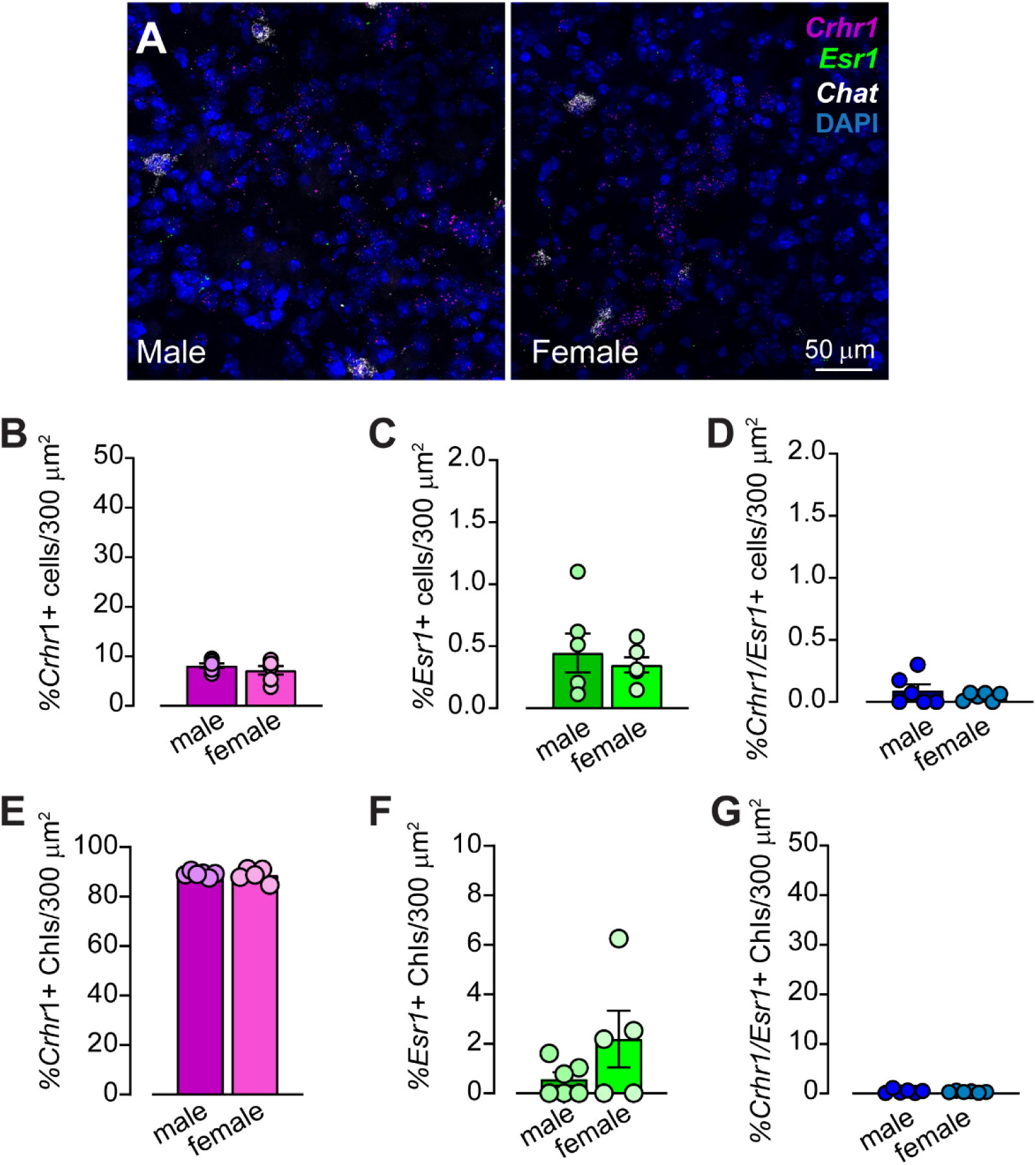
No sex differences in *Crhr1+* and *Esr1+* mRNA in dorsal striatum. A) Example images of *in situ* hybridization showing *Crhr1* (magenta), *Esr1* (green), *Chat* (white), and DAPI (blue) in the male (left) and female (right) dorsal striatum. B) No change in the density of *Crhr1+* cells in the dorsal striatum of males and females (unpaired t-test, t = 0.9380, p = 0.3703, N = 6 males, 6 females). C) The density of *Esr1+* cells in the dorsal striatum of males and females is similarly sparse (unpaired t-test, t = 0.5742, p = 0.5785, N = 6 males, 6 females). D) There is minimal overlap in the expression of *Crhr1* and *Esr1* in the dorsal striatum in male and female mice (unpaired t-test, p = p = 0.5785, N = 6 males, 6 females). E) Percentage of *Chat*+ neurons (putative cholinergic interneurons, ChIs) co-expressing *Crhr1* is similarly high in both male and female mice (unpaired t-test, p = 0.7469). F) Percentage of ChIs co-expressing *Esr1* is relatively low in both male and female mice, with female mice showing an increase in inter-animal variability compared to male mice (F test to compare variances, p = 0.0124), but no change in mean percentage of ChIs expressing *Esr1* (unpaired t-test, p = 0.1666, N = 6 males, 6 females). G) There is virtually no overlap in the expression of *Crhr1* and *Esr1* in ChIs in the dorsal striatum in either males or females (unpaired t-test, p = p >0.9999, N = 6 males, 6 females).

We next tested where there were differences in ChI-specific expression of *Crhr1* or *Esr1* in the dorsal striatum of males and females. As previously reported in males only^19^, *Crhr1* was highly co-expressed in *Chat*+ ChIs (89.1±0.4%, N = 6) in male mice, which was matched in female mice (88.7 ± 1.2%, N = 5, unpaired t-test, t = 0.3328, p = 0.7469, Figure 3a,e). Again, only a very sparse number of ChIs in the dorsal striatum expressed *Esr1* in either males (0.57±0.3%, N = 6) or females (2.2 ± 1.1%, N = 5). While the mean difference in ChI-specific *Esr1* mRNA expression between the sexes trended toward significance (unpaired t-test, p = 0.1666), there was a notably greater variance in ChI-specific *Esr1* mRNA expression in females compared to males (F test to compare variances, p = 0.0124, Figure 3f). Finally, there was negligible co-expression of *Crhr1* and *Esr1* (less than 0.1%) in all cells of the dorsal striatum, and this was true for both males and females (unpaired t-test, t = 0.9035, p = 0.3875, N = 6 males, 6 females, Figure 3d), and the same pattern held in ChIs specifically (unpaired t-test, t = 0.000, p >0.9999, N = 6 males, 6 females, Figure 3g). Taken together, these data show little difference between males and females in either *Crhr1*+ or *Esr1*+ cell quantity and distribution in the dorsal striatum and suggest there is likely no interaction between these two receptors at the cellular level as they are rarely co-expressed. In contrast, as we explored the expression of *Crhr1 and Esr1* in the nucleus accumbens, we uncovered robust sex differences.

### Sex differences in Crhr1 and Esr1 in the NAc

To test if there were sex differences in overall or ChI-specific *Crhr1* or *Esr1* mRNA expression in NAc we used *in situ* hybridization strategies, as above. We found that females had a significantly higher density of *Crhr1*+ cells in the NAc compared to males (intact male: 8.4 ± 0.6%, intact female: 18.1 ± 1.1%, N = 7 males, 29 females, Figure 4a,b). To determine whether this sex difference was driven by endogenously cycling sex hormones, we further analyzed *Crhr1* expression in females that had been ovariectomized (OVX) and then treated with a single administration of 17β-estradiol (E2) or vehicle prior to tissue collection. OVX females had an intermediate level of *Crhr1* mRNA expression and as a result were not significantly different from intact males or intact females. Acute estradiol treatment in OVX females did not restore the higher *Crhr1+* cell density seen in intact females (OVX + VEH female: 12.7 ± 0.8%, OVX + E2 female: 13.6 ± 1.1%, one-way ANOVA, F_3,41_ = 9.704, p < 0.0001, Tukey’s post-hoc t-test, intact male vs intact female, p < 0.0001, intact female vs. OVX + VEH, p = 0.1328, intact female vs. OVX + E2, p = 0.0791, intact male vs OVX + VEH, p = 0.4240, intact male vs OVX + E2, p = 0.1288, N = 4–29, Figure 4a,b). Although ovariectomy obscured the sex difference in *Crhr1*+ cell density, we did not find any differences in intact females across the estrous cycle (one-way ANOVA, F_3,21_ = 1.463, p = 0.2518, N = 4-9, Figure 4c).

**Figure 4.**
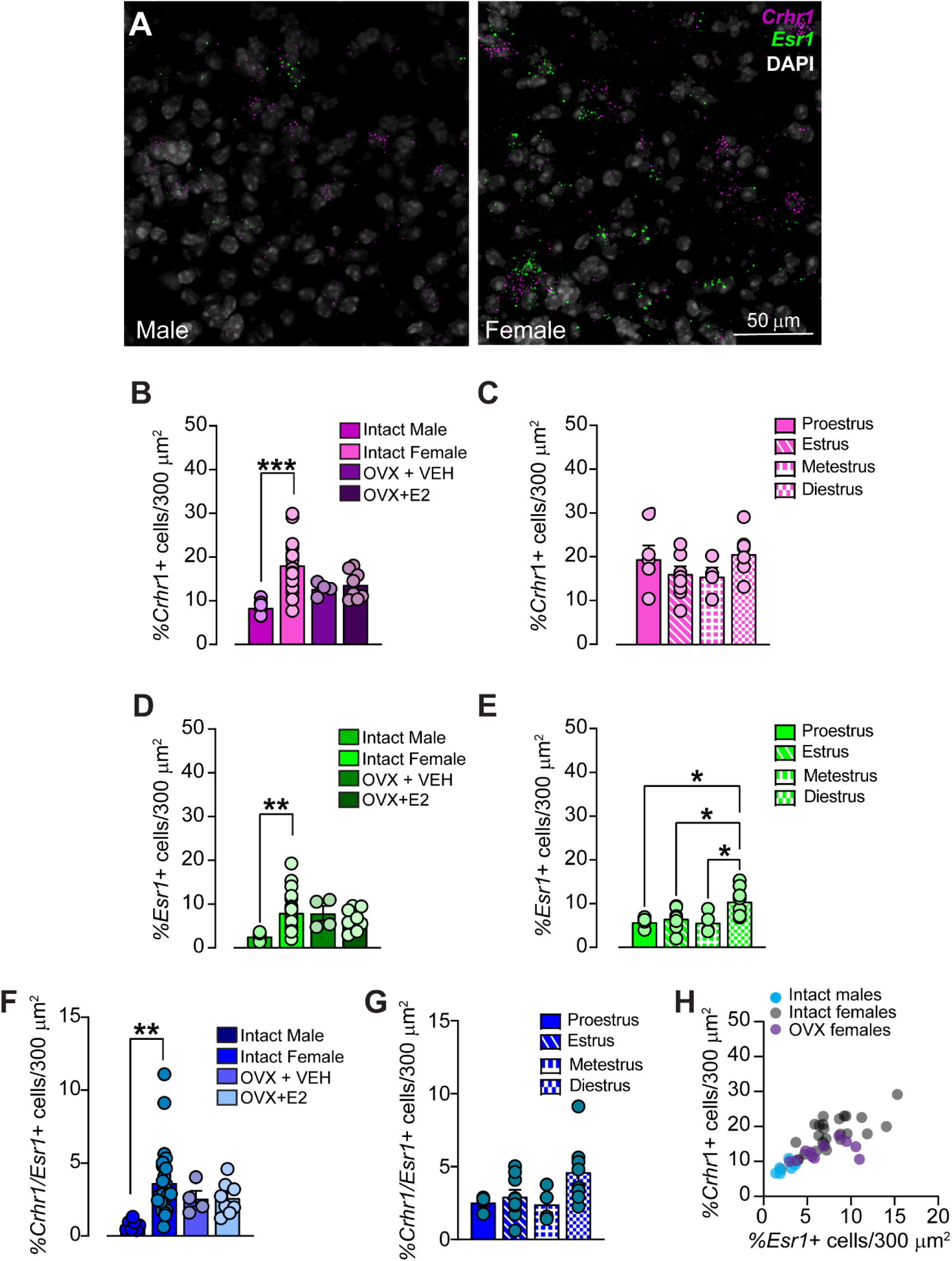
Sex-dependent differences in Crhr1+ and Esr1+ mRNA in the nucleus accumbens (NAc). A) *In situ* hybridization showing *Crhr1* mRNA (magenta), *Esr1* mRNA (green), and DAPI (white) in male (left) and female (right) NAc. B) The density of *Crhr1+* cells is higher in the NAc of females than males (one-way ANOVA, F_3,41_ = 9.704, p < 0.0001, N = 4–29), with intermediate *Crhr1+* cell densities seen in ovariectomized (OVX) females after acute administration of either vehicle (VEH) or 17β-estradiol (E2). C) The density of *Crhr1+* cells does not change across phases of the estrous cycle in female mice (one-way ANOVA, F_3,21_ = 1.463, p = 0.2518, N = 4-9). D) The density of *Esr1+* cells is higher in the NAc of females than males, with *Esr1+* cellular densities maintained in OVX females treated with either VEH or E2 compared to intact females (one-way ANOVA, F_3,43_ = 5.073, p = 0.0044, N = 4– 29). E) *Esr1+* cell density fluctuates over the estrous cycle in intact females, with the highest percentage of *Esr1+* cells found during diestrus (one-way ANOVA, F_3,21_ = 5.560, p = 0.0057, N = 4–9). F) Females have a higher number of cells that co-express *Crhr1* and *Esr1* in the NAc compared to males, with OVX females showing an intermediate phenotype (one-way ANOVA, F_3,41_ = 4.490, p = 0.0082, N = 4–29). G) The density of *Crhr1+/Esr1+* cells does not change across phases of the estrous cycle in female mice (one-way ANOVA, F_3,21_ = 2.694, p = 0.0721, N = 4-9) H) There is a positive correlation between the density of *Crhr1+* cells and *Esr1+* cells in all mice regardless of sex or cycling hormones (intact mice: Pearson’s r = 0.72, p < 0.0001; OVX mice: Pearson’s r = 0.577, p = 0.049). *: Tukey’s post-hoc t-tests, p < 0.05, **: Tukey’s post-hoc t-test, p < 0.01 ***: Tukey’s post-hoc t-test, p < 0.0001

We next assessed whether there were changes in *Esr1* mRNA expression in the NAc of males and females. Similar to the pattern we found in *Crhr1* expression, we observed a significant increase in *Esr1*+ cells in the NAc of females compared to males, though this was not significantly reversed in OVX animals administered vehicle or E2 (intact male: 2.6 ± 0.3%, intact female: 8.0 ± 0.8%, OVX + VEH female: 7.9 ± 1.6%, OVX + E2 female: 6.6 ± 0.9%, one-way ANOVA, F_3,43_ = 5.073, p = 0.0044, Tukey’s post-hoc t-test, male vs female, p = 0.0023, male vs OVX + VEH female, p = 0.0645, male vs OVX + E2 female, p = 0.1111, N = 4–29, Figure 4d). Unlike with *Crhr1*+ cells, however, we did find a difference in density of *Esr1+* cells across the estrous cycle, with diestrus females showing a higher density of *Esr1*+ cells compared to proestrus, estrus and metestrus females (diestrus: 10.5 ± 1.1%, proestrus: 5.8± 0.6%, estrus: 6.6 ± 0.8%, metestrus: 5.7 ± 1.2%, one-way ANOVA, F_3,21_ = 5.560, p = 0.0057, Tukey’s post-hoc t-tests, p < 0.05, N = 4–9, Figure 4e). There was a small percentage of NAc cells that co-expressed *Crhr1* and *Esr1* and similarly to what we found with the individual mRNA analysis, females had a higher co-expression than males, with OVX females showing an intermediate level between intact males and females (intact male: 0.74 ± 0.13%, intact female: 3.6 ± 0.45%, OVX + VEH female: 2.6 ± 0.53%, OVX + E2 female: 2.6 ± 0.40%, one-way ANOVA, F_3,41_ = 4.490, p = 0.0082, Tukey’s post-hoc t-test, male vs female, p = 0.0045, male vs OVX + VEH female, p = 0.4491, male vs OVX + E2 female, p = 0.2397, N = 4–29, Figure 4f). There were no significant differences across estrous, though there was a trend for a similar pattern to what was seen with the individual *Esr1* analysis (one-way ANOVA, F_3,21_ = 2.694, p = 0.0721, N = 4-9). Finally, we found a significant positive correlation between the density of *Crhr1+* and *Esr1+* cells in the NAc in intact males and females (Pearson r correlation = 0.72, p < 0.0001, Figure 4h). The relationship between *Esr1* and *Crhr1* mRNA in ovariectomized females was also significant (Pearson r correlation = 0.577, p = 0.049). In summary, we found that overall expression of both *Crhr1* and *Esr1* was higher in females, with *Crhr1* expression sensitive to ovariectomy but not the estrous cycle, and *Esr1* expression sensitive to estrous cycle but not ovariectomy. Further, our correlational data suggests a positive relationship between the expression of these two receptors that is insensitive to the presence of cycling sex hormones.

### Sex differences in Crhr1 and Esr1 expression in NAc ChIs

We next analyzed the expression and co-expression of *Crhr1* and *Esr1* on *Chat*+ NAc ChIs in the same group of mice to determine if the patterns we found in overall *Crhr1* and *Esr1* expression would be mirrored in ChIs. To our surprise, we found intact females had a lower level of *Crhr1* expression in ChIs compared to intact males and this was reversed in OVX females and not recovered by acute E2 injection (intact male: 81.2 ± 3.4%, intact female: 69.8 ± 2.0%, OVX + VEH female: 91.3 ± 0.6%, OVX + E2 female: 91.1 ± 1.0%, one-way ANOVA, F_3,44_ = 16.03, p < 0.0001, Tukey’s post-hoc t-test, male vs female, p = 0.0259, female vs. OVX + VEH, p < 0.0001, female vs. OVX + E2, p = 0.0005, N = 4–29, Figure 5a,b). Additionally, there was a significant effect of estrous cycle: females in estrus had higher levels of *Crhr1/Chat* co-expression compared to females in diestrus (proestrus: 70.12± 3.5%, estrus: 76.3 ± 1.7%, metestrus: 71.7 ± 6.8%, diestrus: 61.5 ± 4.4%, one-way ANOVA, F_3,22_ = 3.269, p = 0.0405, Tukey’s post-hoc t-test, estrus vs. diestrus, p = 0.0249, N = 4–9, Figure 5c). These two findings were the reverse of the patterns we observed across all neurons in the NAc (Figure 4b,c).

**Figure 5.**
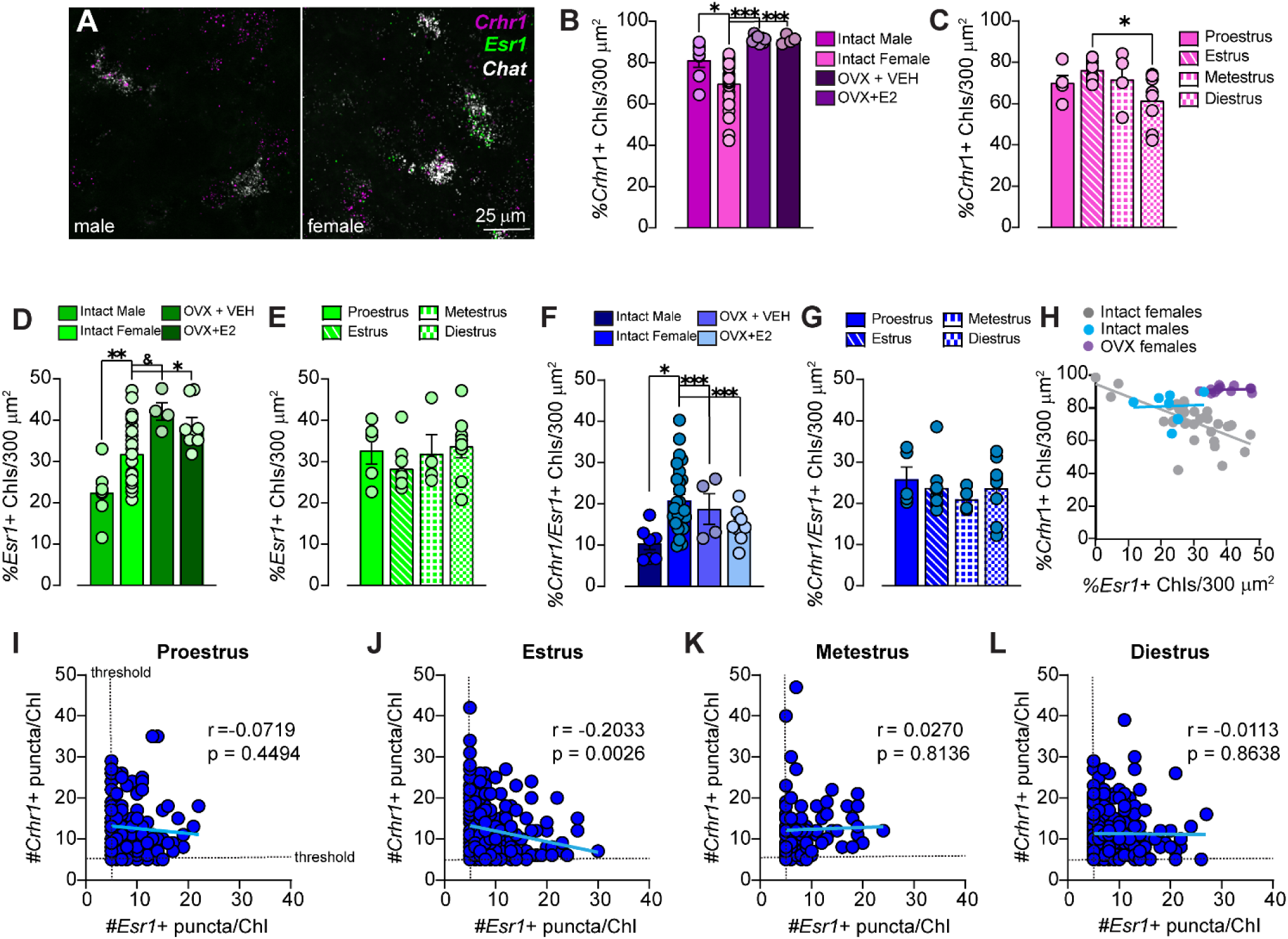
Sex-dependent differences in *Crhr1+* and *Esr1+* co-expression in nucleus accumbens (NAc) cholinergic interneurons (ChIs). A) Example images of *in situ* hybridization showing *Crhr1* mRNA (magenta), *Esr1* mRNA (green), and *Chat* mRNA (grey, putative ChIs) in male (left) and female (right) NAc. B) Fewer ChIs co-express *Crhr1* mRNA in intact females compared to intact males or ovariectomized (OVX) females treated acutely with vehicle (VEH) or 17β-estradiol (E2) (one-way ANOVA, F_3,44_ = 16.03, p < 0.0001, N = 4–29). C) NAc ChIs express *Crhr1* mRNA across the estrous cycle in female mice, with a dip in *Crhr1* mRNA expression during diestrus compared to estrus (one-way ANOVA, F_3,22_ = 3.269, N = 4–9). D) Fewer ChIs co-express *Esr1* mRNA in intact males compared to intact females or OVX females treated with either VEH or E2 (one-way ANOVA, F_3,44_ = 10.47, p < 0.0001, N = 4–29). E) *Esr1* mRNA expression in ChIs does not fluctuate across the estrous cycle in intact females (one-way ANOVA, F_3,22_ = 0.8786, p = 0.4673, N = 4–9). F) Co-expression of *Crhr1* with *Esr1* mRNA in NAc ChIs is higher in intact females compared to intact males or OVX females treated with either VEH or E2 (one-way ANOVA, F_3,44_ = 24.72, p < 0.0001, N = 4–29). G) Co-expression of *Crhr1* with *Esr1* mRNA in NAc ChIs does not fluctuate across the estrous cycle in intact females (one-way ANOVA, F_3,22_ = 0.4508, p = 0.7193, N = 4–9). H) There is a strong negative correlation between the percentage of ChIs expressing *Crhr1* mRNA and the percentage of ChIs expressing *Esr1* mRNA in intact female mice (Pearson’s r = −0.63, p < 0.0001), which is not in OVX females (Pearson’s r = 0.12, p > 0.05) or intact males (Pearson’s r = 0.05157, p = 0.9126). I-L) The number of *Crhr1* mRNA puncta seen per ChI is not correlated with the number of *Esr1* mRNA puncta per ChI during proetrus (I), metestrus (K), or diestrus (L) but is negatively correlated with *Esr1* mRNA puncta per ChI during the estrus phase (J) (Pearson’s r = −-0.2033, p > 0.0026). *: Tukey’s post-hoc t-test, p < 0.05 **: Tukey’s post-hoc t-test, p < 0.01 ***: Tukey’s post-hoc t-test, p < 0.0001 &: Tukey’s post-hoc t-test, p < 0.1

We next assessed the expression of *Esr1* mRNA expression in ChIs. *Esr1* mRNA was expressed in a greater percentage of ChIs compared to all NAc cells (compare Figure 5d to Figure 4d, for example). However, the pattern of sex- and hormone-related differences were similar. Females had higher expression of *Esr1* mRNA in ChIs compared to males, and this was observed in both intact and OVX females. In fact, ovariectomy led to a trend in even higher levels of ChI-specific *Esr1* mRNA levels, although with E2 treatment this was reduced to only a trend compared to intact females (intact male: 22.4 ± 2.4%, intact female: 31.8 ± 1.3%, OVX + VEH female: 42.1 ± 2.1%, OVX + E2 female: 38.6 ± 2.0%, one-way ANOVA, F_3,44_ = 10.47, p < 0.0001, Tukey’s post-hoc t-test, male vs female, p = 0.0087, female vs. OVX + VEH, p = 0.0286, female vs. OVX + E2, p = 0.0657, N = 4–29, Figure 5d). However, unlike total *Esr1+* cell number, we did not see any effect of estrous cycle on *Esr1* mRNA levels in ChIs specifically (Figure 5e). We identified similar patterns when examining *Crhr1/Esr1* co-expression in ChIs (Figure 5f,g). Females had a higher frequency of *Crhr1/Esr1* mRNA co-expression in ChIs despite having fewer *Crhr1+* ChIs, and this was observed in both intact and OVX females (intact male: 14.89 ± 2.4%, intact female: 23.2 ± 1.1%, OVX + VEH female: 38.57 ± 2.3%, OVX + E2 female: 36.1 ± 2.0% one-way ANOVA, F_3,44_ = 24.72, p < 0.0001, Tukey’s post-hoc t-test, male vs female, p = 0.0082, female vs. OVX + VEH, p < 0.0001, female vs. OVX + E2, p < 0.0001, N = 4–29, Figure 5f). When assessing whether ChI-specific co-expression of *Crhr1* and *Esr1* changed across the estrous cycle, we found that, like *Esr1* expression but unlike *Crhr1* expression, there were no effects of estrous cycle on the co-expression of *Crhr1* and *Esr1* mRNA in ChIs (Figure 5g). Finally, we found a significant negative correlation between *Crhr1* and *Esr1* in ChIs intact females (Pearson r correlation = −0.44, p < 0.0001, Figure 5h), as opposed to the significant positive correlation we observed in all NAc cells. Interestingly, this relationship was flattened in intact males (Pearson r correlation = 0.0026959, p = 0.9156, Figure 5h). Likewise, ovariectomy flattened this relationship (Pearson r correlation = 0.12, p > 0.05, Figure 5h). Collectively, these data suggest a unique pattern of ERα and CRF-R1 receptor co-expression on ChIs compared to other cell types in the NAc.

To achieve a more granular understanding of the relationship between *Esr1* and *Crhr1* expression, we next quantified the puncta detected by in situ hybridization for each gene in NAc ChIs that co-expressed both receptors’ mRNA in female mice across the estrous cycle. The number of puncta detected by in situ hybridization is directly proportional to the number of mRNA transcripts present in the cell. We found a significant negative correlation between *Esr1* mRNA puncta and *Crhr1* puncta only when females were in estrus and not during other phases (Pearson r correlation = - 0.2033, p > 0.0026, Figure 5i-l). We chose to focus this analysis on individual cells rather than averages across animals because it better captures the variance in puncta number across the population than using animals as the sample number. One caveat to this analysis was that we only examined cells that met our criterion of 5 puncta present to be considered a positive cell. These data provide further evidence for an antagonistic interaction between expression of ERα and CRF-R1 at the cellular level in NAc ChIs.

## Discussion

ChIs are master regulars of striatal output and recent work suggests potential region- and sex-dependent differences in cholinergic regulation of dopaminergic transmission^29,54^. As the number and density of ChIs could impact downstream modulation, we first analyzed if there were any gross differences in ChI numbers across region or sex. We replicated our results and that of others showing that the NAc core has a significantly lower number of ChI somata compared to areas of the dorsal striatum or NAc shell^19^. Yet, we found mixed results when looking at sex differences in ChAT protein and RNA levels. While we did not detect sex differences in number or density of cells immunoreactive for ChAT protein, we did find an increase in the number and density of cells positive for *Chat* mRNA in females. The reasons for this could be technical and/or biological and remain hotly debated^55^. While measures of protein levels are arguably more functionally relevant to the final behavioral output of the animal, the immunohistochemistry method (or any use of antibodies in general such as in western blot) are limited by lower sensitivity. Measures of mRNA tend to be more sensitive, yet the functional relevance of measuring mRNA is unclear^55^. The discrepancy could also be biological. There could be less efficient translation of *Chat* mRNA to ChAT protein in females compared to males. In contrast, there could be more cells capable of releasing acetylcholine in females that are simply not detected using immunohistochemical methods. The next step is to measure acetylcholine release itself. This is an important direction for future research and could be answered using a combination of optogenetic, fiber photometry and electrophysiological techniques.

Our second aim was to assess expression and regulation of CRF-R1 and ERα mRNA in the dorsal and ventral striatum in male and female mice. The antibodies directed towards these receptors, particularly CRF-R1, are not reliable in our experience, necessitating the exclusive used of in situ hybridization methods. Activation of either CRF-R1 or ERα receptors strongly alters activity patterns in the striatum^46,49^, so these receptors are therefore likely to be key players in maintaining (or disrupting) striatal function. Furthermore, many striatal-dependent neurological and neuropsychiatric disorders have different prevalence and even present themselves differently in males and females. Moreover, many of these striatal-linked disorders and their accompanying sex biases including autism spectrum disorder, depression, addiction, ADHD, PTSD, obsessive-compulsive disorders have been linked independently to differences in estrogen and CRF signaling in the brain^13,56–59^.

We therefore analyzed the expression levels in male and female mice and determined the impact of hormonal cycle and ovariectomy on expression of these two receptors throughout the striatum. First, we noted several striking differences between the dorsal striatum and NAc. There was sparse ERα receptor expression in the dorsal striatum, with no significant sex difference. There was also virtually no co-expression of *Crhr1* and *Esr1* in the dorsal striatum of male or female mice in general, or in ChIs specifically. This is despite *Crhr1* being expressed in nearly every ChI in the dorsal striatum. In contrast, we found significant sex differences in *Crhr1* expression, as well as *Esr1* expression, in NAc cells generally and in NAc ChIs specifically.

We were surprised to find differences in the expression patterns of *Crhr1* and *Esr1* when looking at all cells compared to ChIs specifically. For example, when looking for changes in either *Esr1* or *Crhr1* mRNA across estrous, we found differences in cycle-dependent changes depending on whether we were examining all cells or only ChIs (i.e. estrus versus diestrus *Crhr1* expression). Even more surprising was that when looking at all cells, there seems to be a positive correlation between *Crhr1* and *Esr1* that is present in intact and ovariectomized females. In contrast, in ChIs, there is a negative correlation between *Crhr1* and *Esr1* in intact females that is not present in males or ovariectomized females. We can offer some unsatisfying speculations. First, it is possible that the genomic and molecular machinery that CRF-R1 and ERα can interact with is fundamentally different ChIs compared to MSNs, astrocytes or GABAergic interneurons that make up the other cells. It has been shown, for example, that the molecular components that make up the integrated stress response pathway are constitutively active in ChIs compared to other striatal cell types or in most cell types in the brain^60^. It has also been shown that ribosomal phospho-S6 is particularly prevalent and active in ChIs^61^. So, it is possible that ChIs have a very distinct molecular composition aside from that necessary to make acetylcholine that differently governs the interaction of different receptors. What should be most salient, however, is that it is primarily on ChIs where the receptor signaling converges on a cellular level.

Importantly, in the NAc we saw a significant subpopulation of ChIs that co-express *Crhr1* and *Esr1* (2/3 of all *Esr1* ChI expressors). In females we found a negative correlation between the number of CRF-R1 and ERα expressing ChIs. Interestingly, ovariectomy seemed to entirely disrupt this antagonistic relationship; similarly, we did not see a significant correlation in males. Looking at actual puncta number, only in estrus was there a significant negative correlation between CRF-R1 transcripts and ERα transcripts in ChI co-expressors. Together, these data indicate that a protracted disruption in normal hormonal cycling may cause a shift in the interaction between *Crhr1* and *Esr1* expression, without producing appreciable differences in *Crhr1* or *Esr1* expression levels independently.

We believe that this subpopulation of ChIs co-expressing *Crhr1* and *Esr1* represents a unique population within motivation circuits in which stress and sex steroid modulation converge within the same cell. Interestingly, at the level of the hypothalamus, it has been well-documented that stress signaling and associated behavioral responses suppress sexual or reproductive signaling and behavior, and that this is partially driven by CRF suppression of reproductive circuits^62,63^. Based on our findings, it appears that there is a similar antagonist relationship between CRF and estrogen receptor mRNA transcription within ChIs in the NAc. This was somewhat surprising given that acute stressors and reproductive signals can transiently elevate dopamine suggesting some sort of synergism^2,64^. However, previous studies have shown that CRF signaling in the NAc appears to be important for driving exploratory behaviors^68^. We speculate that in periods when females are most sexually receptive (i.e. during estrus) drive to explore the environment is suppressed in order to mate and reproduce successfully. Furthermore, there is evidence that more prolonged stressors can negatively affect accumbal-dependent reproductive and pair-bonding behaviors^65,66^. This relationship may also be more or less prominent with age with a heightened interaction during adolescence and reproductive age and a diminishing interaction during reproductive senescence^67^. This conjecture needs to be formally tested, but it is a possibility given what we know about hypothalamic interactions. We hope that this study is a starting point to explore sex and stress interactions at a mechanistic level within the NAc.

## Acknowledgements

This study was funded by K99/R00 Pathway to Independence award (MH109627) to JCL and NIMH BRAINS R01 (MH122749). Dr. Kavya Devarakonda and Dr. Zoe Christenson-Wick contributed key edits to the manuscript. The authors declare no competing financial interests.

